# Exploring the ageing and survival costs of investment in anti-predation responses in a wild insect?

**DOI:** 10.64898/2026.04.28.721323

**Authors:** Ruonan Li, Rolando Rodríguez-Muñoz, Tom Tregenza, Lucy Winder

## Abstract

Escape behaviour directly influences survival, yet individuals often vary substantially in escape performance. Laboratory studies have documented trade-offs between anti-predator responses and life-history traits, but it remains unclear whether such trade-offs occur under natural predation risk. We studied a natural population of the field cricket *Gryllus campestris*. Mortality risk and behavioural performance are known to change with age in this species. We aimed to determine whether individuals expressing a higher escape response pay a cost in terms of a faster increase in mortality risk with age or a shorter lifespan. We quantified escape speed in response to a vibrational predation cue. We found no clear evidence for a trade-off between escape performance and lifespan or age-specific mortality risk. The relationship between escape speed and the among-individual effect of age differed between sexes: older males showed faster escape speeds compared with younger males, whereas younger females were faster than older females. This pattern is consistent with sex-specific selective disappearance. Individual baseline mortality risk varied with sex and escape speed, but age-dependent mortality did not. It suggests that such trade-offs in the wild may be context- or condition-dependent rather than reflecting a universal life-history trade-off.

## Introduction

Escape behaviour is one of the most widespread anti-predation strategies and is found across animal taxa (Binazzi et al., 2011; Lagos, 2017; Turesson et al., 2009). The ability to detect and respond rapidly to threats directly impacts an animal’s survival probability, and is therefore likely to be under strong selection (Lind & Cresswell, 2005; Stoks et al., 2003).

Nevertheless, it is common to observe substantial variation in escape responses within species (Domenici, 2010; Li et al., 2025; Marras et al., 2011; Winter et al., 2023), raising the question of how this variation can persist. As anti-predator defences are energetically demanding or costly to produce, one possibility is that investment in anti-predator defences is traded-off against other aspects of an individual’s life, including life history traits such as longevity and fecundity. As such, the benefits of expressing escape behaviours are balanced by their costs.

There is broad evidence that morphological anti-predator defences, such as armour, shell thickening, helmets, spines and disruptive coloration, carry life-history costs, consistent with resource allocation trade-offs (e.g., Bourdeau & Johansson, 2012; Hoverman & Relyea, 2009; Kikuchi et al., 2023; Octorina et al., 2022). It has also been shown that predation levels alter life-history strategies: guppy populations in lower river reaches, exposed to higher predation, evolve faster life histories than those upstream. (Stephenson et al., 2015).

However, evidence for trade-offs between life-history parameters and anti-predator traits that are behavioural rather than morphological are limited, likely because it is more difficult to quantify investment in behaviour. Relationships between behaviour and life-history traits are often discussed using the framework of a pace-of-life syndrome (POLS; Réale et al. (2010)), which predicts covariation between behaviour, physiology and lifespan. However, meta-analyses of pace-of-life predictions report little or inconsistent overall support for stable behaviour vs life-history covariation, suggesting that these relationships may not generalise across ecological contexts (Royauté et al., 2018). An important next step is to investigate the trade-off between behavioural responses to threats and life-history traits such as survival and mortality rate under natural conditions. This is particularly important in wild insects, which often experience high predation and represent the most diverse animal taxon on earth.

Increasing exertion requires an increase in metabolic activity, which inevitably incurs costs such as oxidative damage or consumption of energy that could otherwise be used for somatic maintenance, potentially reducing survival probability or lifespan (Monaghan et al., 2009; Speakman et al., 2015). Individuals that invest heavily in escape behaviour may therefore experience faster rates of ageing. More specifically, such individuals may exhibit a low baseline mortality risk if rapid escape reduces predation early in life, but a steeper increase in mortality risk with age as maintenance costs accumulate. In contrast, individuals that invest less in escape behaviour may show a higher baseline mortality risk but a slower increase in mortality with age, reflecting alternative life-history strategies.

We know relatively little about how behavioural performance varies with age in wild insects, and how this impacts survival probabilities (Zajitschek et al., 2020). Locomotor capacity and sensory responsiveness, as key components of predator escape, are often reported to decline with age. For example, a study of multiple species (*Caenorhabditis elegans, Mus domesticus, Canis familiaris, Equus caballus, and Homo sapiens*) identified a pattern of an increase in locomotor performance during development and a decline in old age that was similar across this diverse group (Marck et al., 2017). Similarly, a review in insects illustrated that increasing age is generally associated with locomotor deficits caused by degeneration in the nervous and musculoskeletal systems (Ridgel & Ritzmann, 2005).

As behavioural and physiological differences between the sexes are common across species, it is possible that there are age-related changes in anti-predator behaviour between the sexes. For instance, a study of the field cricket *Gryllus campestris* showed stronger age-related declines in movement in males than in females (Makai et al., 2020). Sex differences in behaviour are often driven by differences in reproductive strategies, with males typically investing more in mate searching and activity, whereas females invest more in survival and future reproduction (Andersson & Iwasa, 1996). This may arise in escape strategies because males experience higher predation risk due to increased movement associated with mate searching. Males may therefore benefit from greater early-life investment in anti-predator responses. Females, in contrast, may adopt more conservative strategies that prioritise maintenance and longer-term survival.

Although age-related physiological changes and their association with behaviour has been widely studied in laboratory insects, especially in *Drosophila* (Promislow et al., 2022), the laboratory environment could generate results that are unrepresentative of natural populations. Even subtle differences in rearing conditions in the laboratory are known to have substantial effects on senescence (Partridge & Gems, 2007) and benign lab environments can mask costs and trade-offs. This issue limits our understanding of how anti-predation traits integrate into life-history strategies in the natural environment. When studying age-related changes in behaviour it is important to account for the fact that cross sectional comparisons may not necessarily reflect intrinsic senescence because individuals that survive longer may not be a random sample of the population (Van de Pol & Verhulst, 2006). It is, therefore, important to account for this potential selective disappearance when attempting to understand whether there are trade-offs between anti-predation responses and mortality rate, while also accounting for the effects of age.

To investigate whether there is a trade-off between anti-predation responses and mortality risk, we studied natural mortality and a conspicuous anti-predator behaviour in wild population of the field cricket (*Gryllus campestris*) in northern Spain. These crickets rely on ground vibrations or air currents to detect approaching predators, and respond by rapidly fleeing into a burrow a few centimetres away (Broder et al., 2021; Li et al., 2025; Stevenson et al., 2000). We simulated predation cues through ground vibrations and measured cricket escape speed (Li et al., 2026). We investigate whether 1) escape performance decreases with age; 2) individuals with faster escape speeds exhibit shorter lifespans or higher mortality rate, consistent with life-history trade-off theory; and 3) mortality risk varies between males and females and their investment in escape speed.

## Methods

### Study system

Our study site is the ‘WildCrickets’ meadow in Gijon, northern Spain where we have been studying a wild population of the field cricket *Gryllus campestris* for many years (Tregenza et al., 2022) (see www.wildcrickets.org). *G. campestris* has annual generations and relies on basking in sunshine to maintain body temperature in the early spring (Alexander, 1968; Gardner et al., 2024; Li et al., 2025; Rodríguez-Muñoz et al., 2023), which exposes them to predation risk. Nymphs excavate burrows from an early age, diapause over winter and emerge in spring, becoming adult in April-May (Rodríguez-Muñoz et al., 2025). Adults are flightless and strongly associated with burrows which they run into very rapidly to escape predators.

In March 2024, we located every burrow in our meadow and marked each of them with a flag carrying a unique number. We monitored 140 of the burrows continuously using infra-red digital video cameras. We began collecting data on each individual only after it became adult. Because adult crickets frequently moved between burrows, leaving one burrow and arriving at another a few minutes later (median duration of each visit to a burrow is 1.2 hr) (Rodríguez-Muñoz et al., 2025). Cameras were periodically relocated from inactive burrows (defined as burrows not used by any crickets for more than two days based on video playback) to active burrows where crickets were observed. This approach allowed us to track the majority of crickets throughout their lifespan within the meadow. Using these observations, we were able to determine an individual’s death by an individuals last sighting or is it based on whether it was observed for 2 consecutive days.

### Escape speed measurement and predation cues stimulus

To simulate the vibrational cues that stimulate crickets to flee down their burrow, we dropped an 40mm diameter balls onto the ground immediately adjacent to the burrow generating a vibrational stimulus. Our previous study used balls of two different masses to generate vibrational stimuli of weak (9 g cork ball) and strong intensity (266 g steel ball) (hereafter referred to as ‘simulation type’) (Li et al., 2026). Each cricket was exposed to both treatments (weak or strong intensity) and the order of the treatments was randomised.

Escape speed assays followed Li et al. (2025). Briefly, a GoPro (240 frames per second, 2.7K pixel resolution) was positioned 50 cm above the burrow. A ruler placed beside the burrow provided scale. When a cricket had remained still outside its burrow for ∼1 min, we recorded its body temperature with an IR thermometer (RS PRO DT-836 or Testo 830-T4), then triggered the video and released the stimulus ball down a 50 cm long tube. Individual escape speed was measured across repeated trials per individual (median =2, range = 1-9) under varying natural environment conditions. We also recorded their age at time of measurement in days from emergence and their sex.

We used the software ‘Kinovea’ (https://www.kinovea.org/download.html) to review videos in slow motion using the frame number displayed with each frame representing 1/240_th_ of a second. Using the ruler as reference, we measured the time from the first detectable movement to the point when the cricket had moved 1.5 cm away from its initial position. Standardising this distance avoided the influence of acceleration which creates differences in average speed among individuals fleeing over varying distances. This 1.5cm distance has been previously validated as a reliable measure for assessing escape speed (Li et al., 2025).

## Data analysis

We measured 91 crickets in total. We excluded 9 individuals due to a lack of information on age or lifespan. The remaining 82 crickets (Female=37, Male=46) provided a total of 253 measurements for analysis (Female=126, Male=127), averaging 3.09 measurements per individual (SD = 1.54).

To examine whether age influences escape performance and whether this relationship differs between the sexes, we modelled escape speed using linear mixed-effects models (LMMs) with Gaussian error distribution implemented in the *lme4* package (v.1.1-36). To address potential selective disappearance (i.e., slower, poor condition individual dying earlier and therefore underrepresented at later measurement windows) (Van de Pol & Verhulst, 2006). We followed Van de Pol & Verhulst’s framework to partition age into within- and among-individual components. Specifically, we calculated within-individual age as the age of an individual at time of observation minus the individual’s mean age across all records. Among-individual age was calculated as each individual’s mean age minus the grand mean of all individuals’ mean ages, giving an estimate of how long a given individual lived for compared to the population mean. Escape speed was included as the response variable, with within-individual age, among-individual age, sex and the interactions between sex and each of the two age components as fixed effects. Individual ID was included as a random effect to account for repeated measures. Body temperature and stimulation type were incorporated as covariates, as both were shown to affect escape speed in previous work (Li et al., 2026; Li et al., 2025). Using this model, we extracted the random intercepts (Best Linear Unbiased Predictors, BLUPs (Robinson, 1991)) to give individual baseline escape speeds for use in further analyses, described below.

To assess whether lifespan varies with escape speed, we fitted a linear model using the *lm* function with lifespan (log-transformed to meet Gaussian error distribution) as the response variable, and baseline escape speed (from BLUPs, as above), sex, and body mass as fixed effects. As mortality hazards can vary with age, we also modelled survival using a Bayesian survival analysis using the ‘*BaSTA’* package (v.2.0.2) (Colchero et al., 2012). We classified individuals into four groups based on sex and baseline escape speed: female-faster, female-slower, male-faster, and male-slower. The faster/slower categories were defined using the median of the overall baseline escape-speed distribution. Survival trajectories of these 4 groups were estimated using BaSTA, with 22,000 iterations, a burn-in of 2,000, and 20 simulations retained, and a ‘simple’ mortality shape within the ‘WE’ (Weibull) model structure was selected. We compared alternative mortality model structures (exponential, Gompertz, Weibull, and logistic) and mortality shapes (simple, Makeham, and bathtub) using the Deviance Information Criterion (DIC). The Weibull model with a simple mortality shape had the lowest DIC and was therefore selected as the best-supported mortality trajectory.

We used the Kullback–Leibler discrepancy calibration (KLDC) to quantify differences in mortality distributions between the four groups of sex and baseline escape speeds, with values > 0.8 indicating substantial differences between groups. As no universal threshold exists for KLDC, we interpret larger values as reflecting greater distributional separation in an information-theoretic sense (Kullback & Leibler, 1951). This approach quantifies pairwise divergence between the posterior distributions for baseline mortality rate (b_0_) or age-dependent mortality rate (b_1_) mortality trajectories, with larger values indicate greater divergence in their estimated b_0_ or b_1_.

All analyses were performed in R v.4.4.2 (R Development Core Team, 2024) via Rstudio 3.03.0 (RStudio Team, 2023). Normality of escape speed was checked using histograms and Q-Q plots, model residuals were inspected to verify the assumptions of homoscedasticity and normality, and multicollinearity was assessed using variance inflation factors (VIFs).

## Results

### 1) Escape speed and contextual factors

As individuals aged, their escape speed tended to increase (within-individual effect), although there was only weak evidence for this pattern (β = 0.013 ± 0.007 SE, p = 0.066; Table 1). We found no evidence of an interaction between within-individual age and sex affecting escape speed (β = 0.009 ± 0.010 SE, p = 0.384; Table 1). There was no among-individual relationship between lifespan and escape speed (β = -0.005 ± 0.004, p = 0.156; Table 1). There was moderate support for variation in escape speed with an interaction between the among-individual effect of age and sex (β = 0.011 ± 0.006 SE, p = 0.048; Table : escape speed was higher in males who on average live longer, but was lower in females who on average lived longer (Figure 1).

**Table 1.** Relationship between escape speed and within-individual age, between-individual age, sex and interaction.

| Predictor | Estimate | SE | df | t value | P value |
| --- | --- | --- | --- | --- | --- |
| Intercept | 0.196 | 0.039 | 229 | 5.006 | **<0.001** |
| Body Temperature | 0.005 | 0.002 | 233 | 2.924 | **0.004** |
| Stimulation (Strong) | -0.042 | 0.012 | 177 | 3.558 | **<0.001** |
| Sex (male) | 0.051 | 0.018 | 86 | 2.875 | **0.005** |
| Age_within | 0.013 | 0.007 | 173 | 1.853 | 0.066 |
| Age_between | -0.005 | 0.004 | 78 | 1.431 | 0.156 |
| Age_within:Sex(male) | 0.009 | 0.010 | 174 | 0.873 | 0.384 |
| Sex(male):Age_between | 0.011 | 0.006 | 87 | 2.008 | **0.048** |

**Figure 1.**
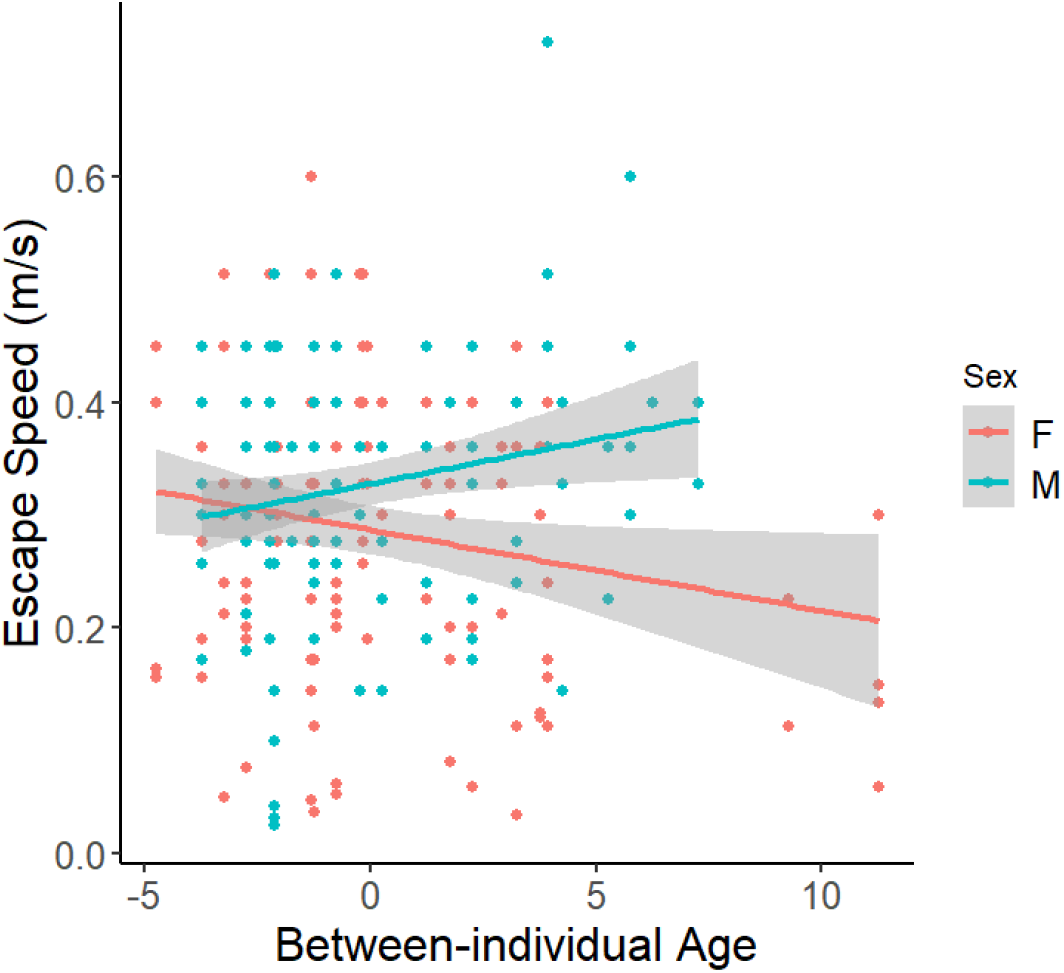
Association of between-individual age with escape speed, stratified by sex. Between-individual age represents each individual’s mean age relative to the grand mean age of the population. Values below zero indicate individuals whose mean age was lower than the population-level mean age. Fitted lines (blue for males, red for females) are displayed with their corresponding data points and a 95% confidence interval (grey shading).

**Figure 2.**
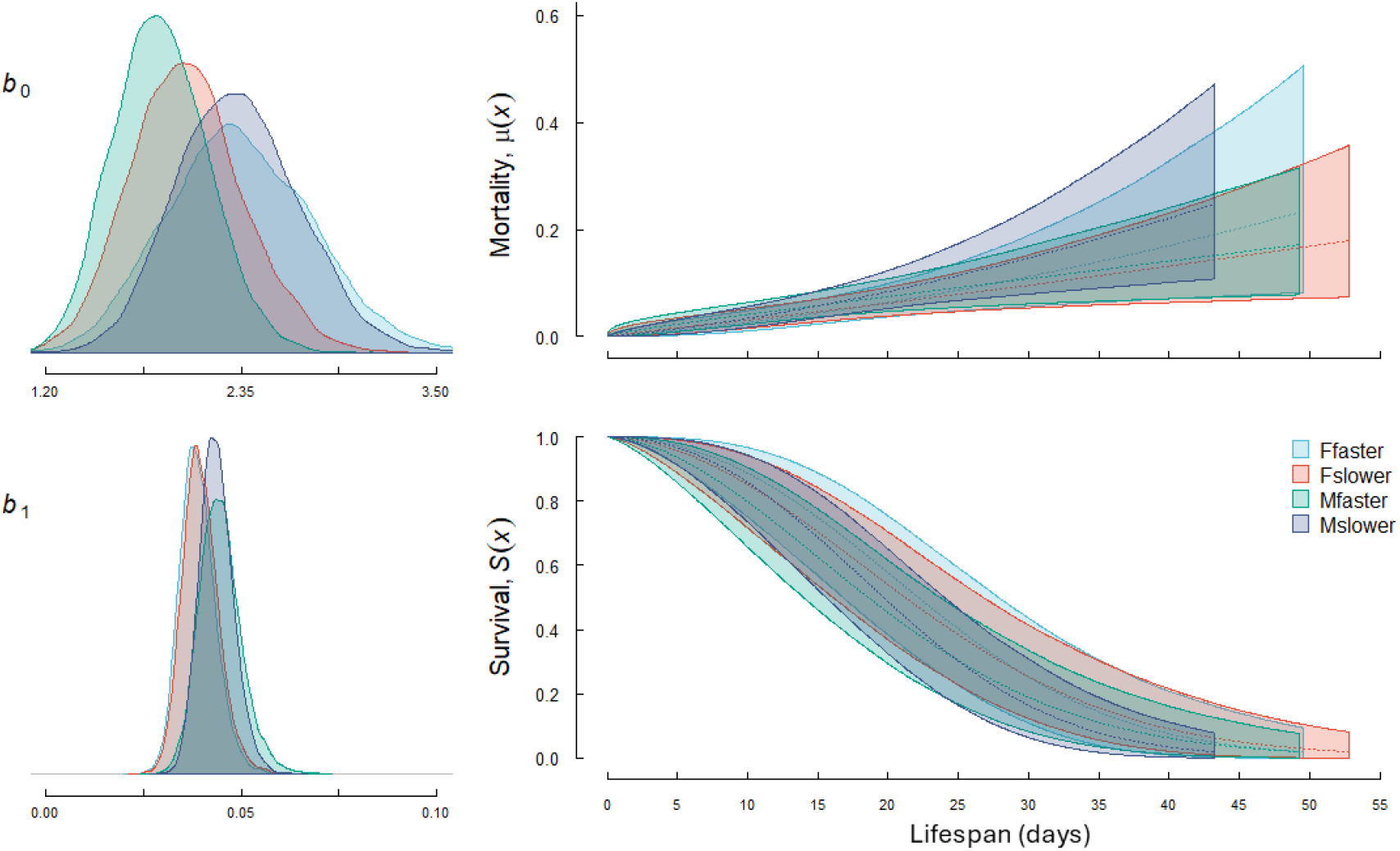
Results from the BaSTA analysis showing estimated mortality and survival trajectories for individuals in 4 (sex × speed) groups. *b*_*0*_ represents the baseline mortality rate and *b*_*1*_ represents the age-dependent mortality rate. Blue and red lines indicate the female-faster and female-slower groups (with 95% CI shaded), respectively. Green and purple lines indicate the male-faster and male-slower groups (with 95% CI shaded), respectively. The two left panels show the posterior distributions for the 4 groups’ baseline mortality (b_0_) and age-dependent mortality rate (b_1_), while the right panels illustrate comparable mortality trajectories across lifespan. Note that parameter estimates for b_0_ are shown on the log scale in the posterior distributions.

In addition, escape speed increased with body temperature (β = 0.005 ± 0.002 SE, p = 0.004; Table 1) and was lower with the strong stimulation type (strong: 0.285 ± 0.011; weak: 0.327 ± 0.011; β = -0.042 ± 0.012 SE, p < 0.001; Table 1). Males had faster escape speeds than females (males: 0.332 ± 0.012 SE; females: 0.281 ± 0.013 SE; β = 0.051 ± 0.018 SE, p = 0.005; Table 1).

### 2) Lifespan and estimated baseline escape speed

We found baseline escape speed did not predict lifespan. Males had shorter lifespans than females (males: 15.9 ± 0.758 SE; females: 19.3 ± 0.925 SE; β = -0.194 ± 0.068 SE, p = 0.0047; Table 2).

**Table 2.** Relationship between lifespan and escape speed, sex and body mass.

| Predictor | Estimate | SE | t value | P value |
| --- | --- | --- | --- | --- |
| Intercept | 2.673 | 0.221 | 12.114 | <0.001 |
| Sex (male) | -0.194 | 0.068 | 2.856 | 0.0047 |
| Mass | 0.307 | 0.229 | 1.343 | 0.180 |
| Estimated speed (BLUPs) | 0.279 | 1.182 | 0.236 | 0.813 |
| BLUPs × Sex (male) | -0.244 | 1.612 | 0.151 | 0.880 |

### 3) Bayesian mortality trajectory analysis (BaSTA)

We found that posterior estimates of baseline mortality (b_0_) varied among some of the four groups but not others, suggesting baseline mortality in some situations varies with both sex and investment in escape speed. The greatest difference in baseline mortality was between fast males (estimate = 1.881 ± 0.281, Table 3) and fast females (estimate = 2.357 ± 0.42, KLDC between groups = 0.835, Table 3). Slow males (estimate = 2.336 ± 0.358, Table 3) also differed significantly in baseline mortality from fast males (estimate = 0.830 ± 0.534, KLDC between groups = 0.830, Table 3). However, baseline mortality between the other groups did not differ significantly (KLDC < 0.8, Table 3). We also found no significant differences in age-dependent mortality (b_1_) among groups (KLDC <0.8, Table 3).

**Table 3.**
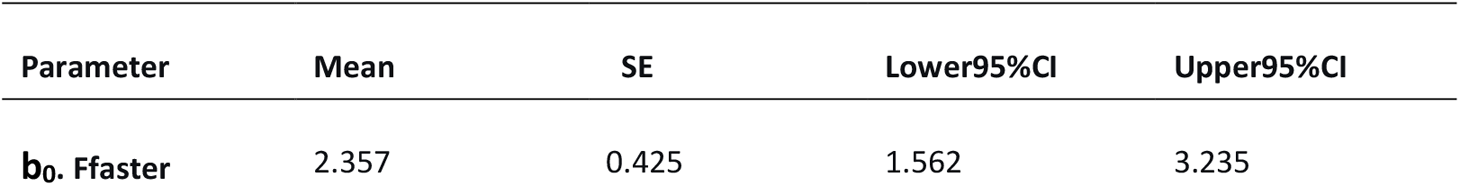
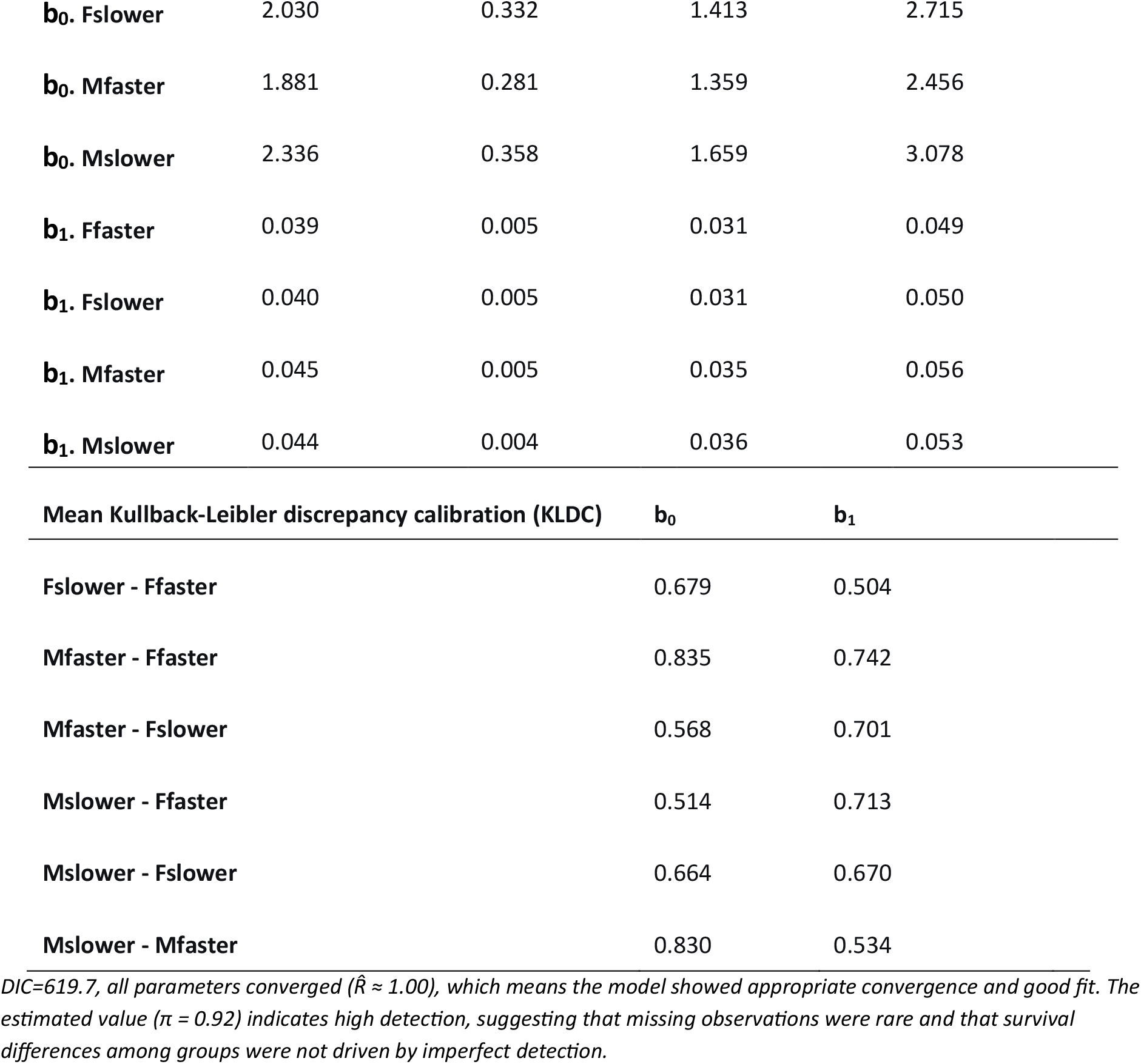
Posterior estimates from Bayesian mortality trajectory analysis (BaSTA). Parameter estimates (mean ± SE) and 95% credible intervals (CI) for Weibull model fitted to 4 (sex × speed) groups.

## Discussion

We tested three main hypotheses in this study: (1) escape performance declines with age; (2) individuals with faster escape speed have shorter lifespans or higher mortality; and (3) mortality risk varies between males and females and their investment in escape speed. Regarding our first hypothesis, we found weak evidence for a positive within-individual age effect on escape speed. This indicates that crickets tend to increase their escape speeds as they age. However, this pattern does not necessarily contradict expectations of senescence, as most crickets in our study were tested between 4 and 21 days after emergence, which represents early to mid-adulthood relative to a maximum lifespan of 49 days in our dataset. This restricted age range means that our data primarily capture early-to mid-life performance and may therefore explain the absence of a detectable decline in escape speed at later life stages. This pattern is consistent with previous work on phenotypic senescence in this species, where performance traits (e.g., singing) often increase during early adulthood before declining at older ages (around 13 or 19 days after becoming adults) (Rodríguez-Muñoz et al., 2019). Similar patterns have been observed across a range of species, including reproductive, physiological, and morphological traits (Monaghan et al., 2008; Nussey et al., 2013). For example, a study on red deer (*Cervus elaphus*) showed that maternal performance initially increased with female age, peaking at around 8 years and then declined until death (Nussey et al., 2006). This non-linear change (increasing during early adulthood, reach a peak, and decline at older ages) suggests that early-life improvement followed by late-life decline may be a general feature of many performance traits.

If escape speed follows a comparable trajectory, this may have implications for sex differences in performance across age. For instance, in red deer (*Cervus elaphus*), males and females begin senescence in survival at similar ages but differ markedly in the rate of senescence (Catchpole et al., 2004). Similarly, in European badgers (*Meles meles*) males exhibit steeper declines in body mass with age than females, a pattern attributed to sex-specific consequences of intense intra-sexual competition experienced during early adulthood (Beirne et al., 2015). More broadly, sex differences in behaviour and ageing trajectories have been linked to differences in reproductive strategies and exposure to predation risk across taxa (Bonduriansky et al., 2008; Christe et al., 2006). In crickets, males may first increase in escape performance and only exhibit senescent declines beyond the age range sampled here. Such a decline, potentially leading male performance to converge with that of females in later life-stage, as locomotor performance often increases during early adulthood before declining at older ages (Marck et al., 2017). The sex-specific patterns are consistent with previous findings in *G. campestris* (Makai et al., 2020), where movement behaviour shows stronger age-related declines in males than in females. Consistent with these patterns, males in our study population also exhibited shorter lifespans than females.

By partitioning age into within- and among-individual components we can not only investigate whether escape speeds decline as individuals get older but, crucially, whether there is selective disappearance of slower crickets. We found that among-individual age effects on escape speed differed between the sexes. Males who lived to be older in the population tended to be faster than males who were recorded whilst young and may have not reached an old age, whereas females displayed the opposite pattern. This pattern is consistent with the possibility of sex-specific selective disappearance, whereby individuals with particular escape performance profiles are more likely to survive to older ages and may reflect differences in life-history strategies and energy allocation between males and females. In many systems, males experience stronger selection on traits related to mate acquisition and competition, which can lead to sex differences in both ageing trajectories and selective disappearance. Adult male crickets engage in energetically costly sexually selected activities such as calling, territorial defence, and mate searching, which may favour individuals with higher locomotor capacity. Females, on the other hand, may allocate more energy to egg production and future reproduction, potentially reducing investment in locomotor performance and resulting in relatively slower escape responses (Andersson & Iwasa, 1996; Roff, 2002).

We also examined the potential anti-predation trade-off with survival by considering two complementary components: lifespan and age-dependent mortality risk. Lifespan reflects overall survival outcomes integrated across the entire life history, whereas mortality risk captures how the probability of death changes with age. We found that contrary to our prediction, baseline escape speed did not predict lifespan. One possible explanation is that in natural environments, the benefits of effective anti-predator behaviour may outweigh its potential energetic or physiological costs. Many studies demonstrating costs of defence or performance traits focus on their energetic or physiological consequences, often revealing trade-offs with other life-history components such as reproduction or maintenance (Guerra, 2011; Guerra & Pollack, 2009; Somjee et al., 2018). However, such approaches typically quantify the costs of trait expression in a predator-free lab environment, without explicitly incorporating the survival benefits of effective anti-predator behaviour. Wild individuals with more effective escape responses may directly reduce their risk of predation and thereby achieve higher survival (Domenici et al., 2011; Lima & Dill, 1990). As a result, underlying trade-offs between investment in escape performance and longevity may be masked or offset by the immediate fitness benefits of predator avoidance. Another possibility is that investment in anti-predator responses affects how mortality risk varies with age, which has important biological implications but cannot be assessed by exploring the effects of anti-predation responses on lifespan alone. As such we also explored age-independent and age-dependent mortality risks separately.

We found some differences among sex-by-speed groups in baseline mortality risk, but these patterns were not consistent across comparisons. In particular, faster females showed higher baseline mortality risk than faster males, and faster males showed lower mortality than slower males (KLDC > 0.8), whereas no clear difference between faster and slower individuals was observed in females. This lack of a consistent pattern suggests that escape speed is not straightforwardly linked to survival benefits across groups. While faster individuals may, in principle, be better able to escape from predators, such benefits do not appear to translate into consistently lower mortality. Instead, variation in baseline mortality may reflect a combination of behavioural, physiological, and condition-dependent factors rather than a simple advantage of faster individuals. In our system, faster crickets may incur energetic or physiological costs, but because escape behaviour is highly plastic and strongly influenced by temperature and cue intensity (Li et al., 2026; Li et al., 2025), much of the observed variation may reflect short-term individual state-dependence or environmental context (Mathot & Frankenhuis, 2018). At the same time, some variation in baseline mortality suggests that escape performance may also partly reflect underlying individual condition. As noted by Van Noordwijk and De Jong (1986), variation in resource acquisition among individuals can obscure trade-offs between life-history traits. In this context, the observed patterns may arise from differences in individual condition (e.g., some individuals can obtain more resources than others) rather than a consistent pace-of-life syndrome.

In contrast, posterior distributions of the age-dependent mortality rate (b_1_) showed substantial overlap and low KLDC values across all four sex-by-speed groups, indicating little evidence that there are differences between groups in their mortality risk with age. This suggests that variation in survival is more likely to arise from differences in baseline mortality rather than in the rate at which mortality increases with age. Together, these results provide no clear evidence for a trade-off between escape performance and mortality risk across sexes. This aligns with meta-analytic evidence showing that associations among pace-of-life traits, including behaviour and life-history characteristics, are often limited and context-dependent across taxa (Royauté et al., 2018). In addition, because most individuals in our dataset were sampled during early to mid-adulthood, any divergence in senescence trajectories may not yet be detectable. Thus, associations between escape performance and mortality may emerge under certain conditions without reflecting a stable life-history correlation among individuals.

Our study also reinforces the strong environmental modulation of anti-predation performance in ectotherms. Escape speed increased with body temperature, as expected from improved muscular performance at higher temperatures (Domenici et al., 2007; Li et al., 2025; Zamora-Camacho et al., 2015). Individuals showed slower escape speed under stronger stimulation, which is consistent with previous findings on anti-predation behaviour in nymphs from the same generation as the adults examined in this study (Li et al., 2026).

Such antipredator performance variations illustrate a context-dependent reaction norm (Mathot & Frankenhuis, 2018; Wolf et al., 2008), and these environmental factors could strongly modulate the trade-offs between behaviour, survival, and lifespan.

Overall, our results suggest that variation in escape behaviour within populations is shaped by both selective disappearance and environmental effects, rather than reflecting intrinsic ageing alone. At the same time, variation in escape performance was strongly associated with environmental conditions, particularly temperature, suggesting that short-term plastic responses may also contribute alongside longer-term life-history processes. Together, these findings indicate that associations between behaviour and survival are context-dependent and may not consistently reflect a stable pace-of-life syndrome. In particular, while we detected differences in baseline mortality among sex-by-speed groups, these patterns were not consistent across all comparisons. Consequently, links between escape performance and mortality appear to depend on both individual condition and environmental context, which may obscure or modify expected life-history trade-offs. More broadly, this suggests that integrating selective disappearance, environmental variation, and phenotypic plasticity is important for understanding how behaviour, survival, and ageing are linked in natural populations.

## Acknowledgments

All authors have no conflict of interests to declare. We sincerely thank Mark Pitt for his help with the data collection in field. This research was supported by a Natural Environment Research Council grant (NE/V000772/1). RL was supported by the China Scholarship Council.

